# Hydrodynamic regime drives flow reversals in suction feeding larval fishes during early ontogeny

**DOI:** 10.1101/771436

**Authors:** Krishnamoorthy Krishnan, Asif Shahriar Nafi, Roi Gurka, Roi Holzman

## Abstract

Fish larvae are the smallest self-sustaining vertebrates. As such, they face multiple challenge that stem from their minute size, and from the hydrodynamic regime in which they dwell. This regime of intermediate Reynolds numbers (*Re*) was shown to affect the swimming of larval fish and impede their ability to capture prey. Numerical simulations indicate that the flow fields external to the mouth in younger larvae result in shallower spatial gradients, limiting the force exerted on the prey. However, observations on feeding larvae suggest that failures in prey capture can also occur during prey transport, although the mechanism causing these failures is unclear. We combine high-speed videography and numerical simulations to investigate the hydrodynamic mechanisms that impede prey transport in larval fishes. Detailed kinematics of the expanding mouth during prey capture by larval *Sparus aurata* were used to parameterize age-specific numerical models of the flows inside the mouth. These models reveal that, for small larvae that slowly expand their mouth, not all the fluid that enters the mouth cavity is expelled through the gills, resulting in flow reversal at the mouth orifice. This efflux at the mouth orifice was highest in the younger ages, but was also high (>8%) in slow strikes produced by larger fish. Our modeling explains the observations of “in-and-out” events in larval fish, where prey enters the mouth but is not swallowed. It further highlights the importance of prey transport as an integral part in determining suction feeding success.

## Introduction

Most marine fish reproduce by broadcasting small (^~^1 mm in diameter) eggs into the open ocean, providing no parental care from the hatching larvae (Blaxter, 1988; Cowen, 2002; Houde, 1987). Typically, larvae deplete their yolk sac after a couple of days (usually 3-7, depending on temperature and environmental conditions) and resort to feed autonomously to gain the necessary resources to complete their development (Blaxter, 1988; Cowen, 2002; Houde, 1987). Despite the staggering variation in body size and life history strategies, the small eggs and larvae, and the lack of parental care, are nearly ubiquitous across marine fish (Barneche et al., 2018). Consequently, fish larvae are the smallest self-sustaining vertebrates. Almost all larval fishes feed in the pelagic realm using “suction feeding”, a characteristic behavior in which fish sequentially open their mouth, expand their buccal cavity and open the opercula covers to generate a unidirectional flow of water that carries their prey into the mouth (Day et al., 2015; Holzman et al., 2015).

In the wild, larval fish suffer dramatic mortality rates (Hjort, 1914; Houde, 1987). It is estimated that >90% of the brood is eradicated during the “critical period”, extending from the time of first feeding until the larvae is ready to settle in its juvenile habitat. During this period, larval fish undergo dramatic morphological and developmental changes, including the ossification of the cranium and vertebrae, the degradation of the fin fold and development of fin rays, as well as the continuous growth and development of the eyes (Blaxter, 1988; Kavanagh and Alford, 2003). Concomitantly, coordination and motor pattern change and improve (Westphal and O’Malley, 2013). The physical growth of the larvae, coupled with the development of stronger muscles that support faster movements lead to an ontogenetic transition in the ways larvae interact with their fluid environment (China and Holzman, 2014; Holzman et al., 2015). Being small and slow, young larvae live in a domain of intermediate Reynolds numbers (*Re*), in which viscous forces are non-negligible compared to inertial ones. This hydrodynamic regime was shown to impede the feeding rates of larval fishes, with 8 Days Post Hatch (DPH) *Sparus Aurata* larvae failing to capture non-evasive prey in ^~^80% of their feeding strikes (China and Holzman, 2014). Manipulations of the viscosity of the medium in which larvae fed demonstrated that the feeding rates of larvae were determined primarily by the hydrodynamic environment, described by the Reynolds numbers that characterized the feeding events (China and Holzman, 2014; Holzman et al., 2015). Older larvae (13 and 23DPH) that fed in a viscous medium displayed feeding rates equivalent to those of the 8DPH larvae in unmanipulated water. Larvae that were raised in mediums with increased viscosity expressed elevated levels of hunger-related neuropeptides (Koch et al., 2018) and suffered higher mortality rates (Yavno and Holzman, 2018). Furthermore, the probability of executing successful prey-acquisition strikes increased with increasing *Re* number calculated for the suction feeding strike (China et al., 2017). Transition into higher Re also improves the larvae’s ability to capture highly evasive prey such as copepods (Jackson and Lenz, 2016; Sommerfeld and Holzman, 2019; Yaniv et al., 2014).

Observations using high-speed videos indicate that one of the reasons for failure in prey acquisition strikes is the occurrence of “in-and-out” events, in which prey is carried into the mouth by the suction flows, but is expelled before the mouth is closed (China et al., 2017; Holzman et al., 2015). The suction flows in these “in-and-out” events were characterized by lower *Re* compared to those in successful events. Furthermore “in-and-out” strikes were initiated from a further distance and were slower compared to unsuccessful events in which the prey did not even enter the mouth (China et al., 2017). A flow visualization study reported flow reversals in larval zebrafish, that occurred in smaller larvae at the time when the mouth started closing (Pekkan et al., 2016). This is in sharp contrast to adult fish, in which flow reversals are rare and minor (Jacobs and Holzman, 2018). However, the extent of these flow reversals across species and developmental stages are unclear, as well as the hydrodynamic conditions under which they occur.

Here, we used computational fluid dynamics (CFD) to investigate the fluid dynamics of suction feeding larval fish. Following Yaniv et al (Yaniv et al., 2014), we constructed a model of an expanding buccal cavity, which incorporates an anterior-to-posterior wave of buccal expansion (Bishop et al., 2008) over time. Our modeling included the opening of the opercula covers at the posterior end of the mouth, a hallmark feature of suction feeding in fishes, which generate unidirectional flows into the mouth while it is closing (Van Wassenbergh, 2015). The model was parametrized based on observed strike kinematics of *Sparus aurata* larvae ranging from first feeding to metamorphosis. Using these kinematics, we quantified the flow speeds and the influx and efflux into the mouth and out of the gills for six larval ages. We then characterized the extent of flow reversals, the flow conditions in which they occur, and the role of hydrodynamics and kinematics (behavior) in driving these flow reversals.

## Methods

### Study organisms

We reanalyzed high-speed videos of suction feeding gilthead sea-bream larvae (*Sparus aurata* Linnaeus, 1758) feeding on Rotifers (*Brachionus rotundiformis;* ^~^0.16 mm in length), from dataset previously used in China and Holzman, 2014 and China et al., 2017. *S. aurata* is a pelagic spawner, hatching at ^~^3.5 mm. Feeding initiates at ^~^5 days post hatching (DPH) at a body length of ^~^4 mm. Larvae reach the stage of flexion at ^~^21-24 DPH, at a length of 7-10 mm, depending on conditions. *Brachionus rotundiformis* is a species of planktonic rotifer, actively swimming at ^~^0.2 mm s^−1^. Prey swimming speed is an order of magnitude slower than the swimming speed of the larvae, and their escape response is considered weak (China and Holzman, 2014; China et al., 2017). Rotifers are universally used as the standard first-feeding food in the mariculture industry.

### High-speed videos

Suction feeding events of larval fish were recorded using high speed video (500 and 1000 frames per second) as described in (China and Holzman, 2014; China et al., 2017). In these experiments, fish swam freely in an aquarium, and their orientation with respect to the camera included lateral, dorsal and ventral views. From the larger dataset of prey acquisition strikes we selected 63 clips in which we could clearly track the kinematics of mouth opening as well as either the hyoid (using lateral view of the fish) or the opercula (using dorsal or ventral views) throughout prey acquisition strike. Clips were selected for fish at the ages of 8, 12-13, 17-18, 22-25, 30 and 35-37 DPH (hereafter 8, 13, 18, 23, 30 and 37 DPH; 4-14 clips per age group). From the lateral view videos, we measured the time of mouth opening and closing, maximal mouth diameter, the time of initiation and peak hyoid displacement and its maximal excursion, and the time of opercula opening and closing (when clearly visible). From the dorsal and the ventral view videos, we measured the time of the mouth opening and closing, the time of initiation and peak opercula displacement and its maximal excursion, and the corresponding parameters at the base of the opercula (1^st^ gill arch). To enable comparisons between different ages and strikes, we standardized the times of hyoid and opercula excursions by the time to peak gape opening (TTPG) in each clip, and their excursions by peak gape. Not all the parameters were visible in all the clips, resulting in a sparse matrix that was ^~^60% full. We averaged the timing and excursion parameters for each landmark, regressed them against larval age and used the predicted values from the regression to generate characteristic kinematics for each age (table 1).

**Table 1:** kinematics characteristics of larval fish used to parametrize the numerical model

| Age (DPH) | Buccal cavity length (mm) | Maximal gape diameter (mm) | Swimming speed (mm/s) | Time to peak gape (s) | Re<sub>L</sub> | inlet |  | outlet |  | *Efflux influx* at inlet (%) |
| --- | --- | --- | --- | --- | --- | --- | --- | --- | --- | --- |
|  |  |  |  |  |  | Peak flow speed (mm/s) | Peak flow rate (mm<sup>3</sup>/s) | Peak flow speed (mm/s) | Peak flow rate (mm<sup>3</sup>/s) |  |
| Observed kinematics |  |  |  |  |  |  |  |  |  |  |
| 8 | 0.81 | 0.11 | 8.1 | 57.3 | 22.8 | 28.3 | 0.58 | 8.50 | 0.13 | 10.1 |
| 13 | 1.06 | 0.15 | 10.6 | 50.1 | 24.4 | 23.1 | 1.37 | 11.1 | 0.35 | 3.1 |
| 18 | 1.23 | 0.17 | 12.3 | 42.9 | 44.1 | 35.9 | 2.24 | 14.3 | 0.61 | 1.3 |
| 23 | 1.36 | 0.19 | 13.6 | 35.7 | 67.6 | 49.8 | 2.96 | 17.3 | 0.93 | 0.34 |
| 30 | 1.50 | 0.21 | 15.0 | 25.7 | 121.3 | 81.3 | 5.22 | 17.4 | 1.12 | 0.26 |
| 37 | 1.61 | 0.23 | 16.1 | 15.6 | 218.2 | 136.2 | 8.36 | 30.9 | 2.26 | 0.06 |
| Low effort strikes |  |  |  |  |  |  |  |  |  |  |
| 23 | 1.36 | 0.19 | 13.6 | 57.3 | 55.2 | 40.7 | 2.6 | 12.6 | 0.62 | 8.2 |
| 30 | 1.50 | 0.21 | 15.0 | 57.3 | 72 | 48.2 | 3.8 | 13.6 | 0.73 | 8.0 |
| 37 | 1.61 | 0.23 | 16.1 | 57.3 | 81.2 | 50.7 | 4.7 | 15.2 | 1.01 | 7.4 |

### Geometry of the modeled buccal cavity

We build on a previous model of mouth cavity expansion suggested by (Bishop et al., 2008; Yaniv et al., 2014), but added the opening of the opercular slits, a hallmark of suction feeding across fishes (Van Wassenbergh, 2015). In brief, the model was composed of three compartments of constant axial lengths, *L*_1_, *L*_2_ and *L*_3_ (Fig. 1). These compartments represented the region from the mouth opening to the anterior hyoid (*L*_1_), the region spanning the anterior to posterior length of the hyoid (*L*_2_) and the region posterior to the hyoid extending to the opening of the esophagus (*L*_3_). Mouth cavity expansion was simulated as time-dependent changes in the radii (, *R*_1_, *R*_2_, *R*_3_, and *R*_4_) of the bases of the compartments, parametrized according to the observed kinematics of the corresponding landmarks in our larvae (see above). The radius *R*_1_ represents the radius of the gape. The lengths *B*_1_, *B*_2_ and *B*_3_ of the lateral surfaces of each compartment varied with time to fit the length variations of the radii *R*_1_, *R*_2_ and *R*_3_. We simulated mouth expansion for six larval ages (8, 13, 18, 23, 30 and 37 DPH) with increasing gape diameter and mouth lengths. Internal dimensions of *L*_1_, *L*_2_ and *L*_3_ were 25%, 30% and 45% of the total mouth cavity length *L*, and mouth radii before mouth expansion were set to 2.5% for *R*_1_ and *R*_4_, and 5% for *R*_2_-*R*_3_ (Yaniv et al., 2014).

**Fig. 1:**
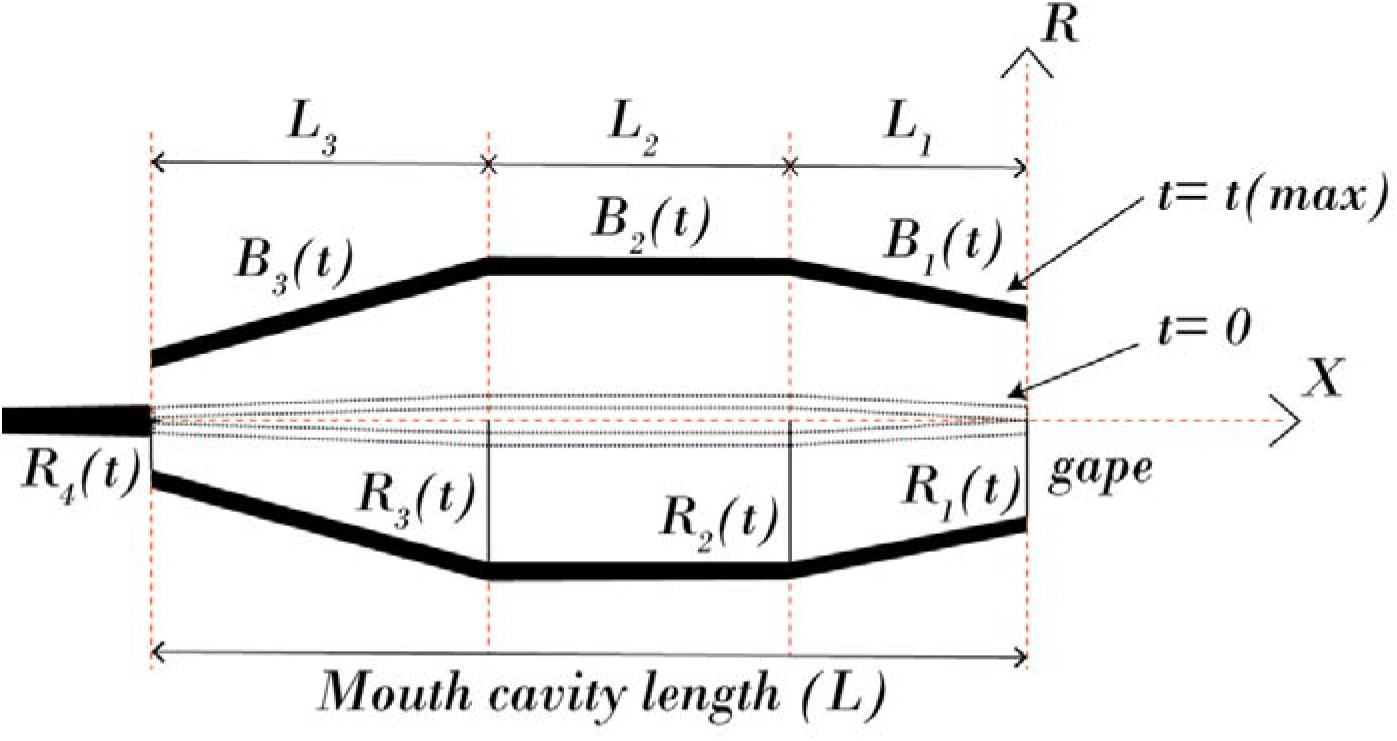
A schematic description of the model geometry. Solid black bars indicate the location of the buccal walls under maximal expansion, light shaded ones show the buccal walls at rest (minimal volume). The mouth is modeled as three attached cones that expend sequentially. L1-L3 correspond to the length of the three cones, whereas R1-R4 is the time-dependent radii of the cones. R1 is the gape (inlet). An increase in R4 represents the opening of the gill slits.

The pattern of mouth opening was simulated by varying the radii *R*(*t*) of each mouth section (*R*_1_-*R*_4_) using the following time-dependent exponential function (Eq. 1; modified from (Müller et al., 1982)):

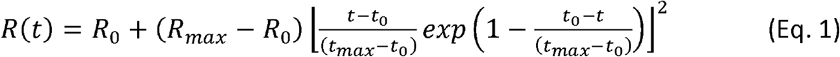

Here, *R*_0_=*R*(*t*=0), *t*_0_ is the time when *R* first deviates from *R*_0_ and *t_max_* is the time when *R* is maximal (*R*=*R*_max_). Note that the radius of each mouth section can have different *R*_0_, *t*_0_, *R*_max_ and *t*_max_ values (Table 1; Fig 3).

Feeding events used to parametrized our model were acquired using manually triggered high-speed cameras, a method which might be biased towards capturing more noticeable events i.e. faster or greater in excursion(China et al., 2017). Thus, they might have represented higher-performance strikes. To investigate the effect of low-effort strikes on the flow dynamics, we run our numerical simulations (see below) for the 23, 30, and 37 DPH cases using the observed geometry but with the expansion kinematics and the relative timing of the 8 DPH case (time to peak gape of 57.3 ms, and time to peak R_2_-R_4_ of 70.3, 73.1 and 78.8 ms, respectively for all three models; Table 1).

### Computational approach

To simulate the fluid dynamics of the buccal cavity and characterize the flow moving in and out of the mouth cavity, a simplified model of an axi-symmetrical mouth cavity was designed. The boundaries of the mouth cavity in the simulations presents a simplified structure that has a cylindrical wall surrounding the cavity and unclosed inlet and outlet edges at the right and left ends, respectively. The cylindrical wall sections are comprised of three length sections that are flexibly connected, and their individual movement was prescribed by the measured kinematics as explained in the kinematics section. To represent the body of the fish and supplement the function of the gills, a streamlined elongated body with a length similar to mouth length was designed downstream the buccal cavity. The body had a small protruding part inside the cavity outlet, with a small (^~^10^−3^ mm) gap from the buccal walls at *t* = 0. At *t* > 0 the mouth started expanding, drawing the fluid in through the gape, followed by the opening of the gap (the gills) based on the prescribed kinematics for R_4_.

The mouth cavity was immersed in a fluid-filled rectangular domain, and it was placed at the center of the domain. The rectangular fluid domain has six boundaries: inlet at the right end, outlet at the left end and four walls (at the top, bottom, far and near) such that uniform flow was formed to move in the domain from right to left. Water at standard atmospheric condition was used as fluid material in the domain. A velocity inlet boundary condition was used at the inlet with water flowing at 13 mm s^−1^ and pressure-outlet boundary condition with standard atmospheric pressure was set at the domain outlet. The top, bottom, far and near walls of the fluid domain, as well as the walls of the mouth cavity were represented with no-slip boundary condition. As the inlet and outlet of the mouth cavity were left unclosed, it is expected for the fluid to flow in and out of the mouth cavity naturally depending on its kinematics. To ensure that the domain size does not interfere with the flow inside and around the mouth cavity, the domain had sufficiently larger dimensions: approximately 30 times the mouth cavity length along the X-direction (flow direction) and 80 times the peak mouth opening radius along the Y-direction for each DPH cases.

The flow field due to expansion of the mouth cavity model was governed by the continuity and momentum conservation equations for incompressible viscous laminar fluid flow in the absence of body force (Ferziger and Peric, 2001). General governing equations for an unsteady, viscous laminar flow is given below.

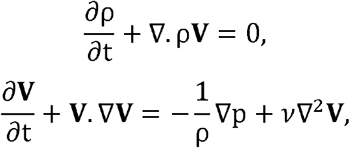

where *ρ* is fluid density, ***V*** is the velocity vector, *p* is the pressure and ν is the kinematic viscosity of the fluid. The flow governing equations were solved using finite volume based commercial software package ANSYS fluent (ANSYS, Canonsburg, Pennsylvania). Mouth cavity model was designed and meshed using ANSYS workbench (ANSYS, Canonsburg, Pennsylvania). Unstructured (triangular shape) mesh method was chosen to discretize the domain, the cavity and its boundaries. Relatively finer meshes were built inside and around the mouth cavity while coarser meshes were used in the domain away from the cavity. To simulate the expansion of the mouth cavity, dynamic mesh method was utilized. The dynamic meshing corresponds to changing of the mesh geometry over time and space based on the prescribed kinematics of the cavity. The kinematic motion of the mouth cavity was prescribed within the fluent solver using the user defined function ‘DEFINE_GRID_MOTION’ (Ansys, 2009; Van Wassenbergh, 2015). This procedure was performed using a user defined function that was compiled and assigned to each length sections of the mouth cavity. Local cell re-meshing method was chosen to re-mesh the mesh grids for every two timesteps based on minimum and maximum cell length and maximum skewness parameters of each cell. To solve the flow equations, a SIMPLE scheme (Ansys, 2009) was employed to carry out the pressure-velocity calculations. Spatial discretization was assigned with second order least square cell-based gradients method whilst a first order implicit method was used for time discretization. The complete numerical solution was obtained by ensuring that the convergence criteria of 10^−4^ for the continuity and the flow speed components are attained. Before proceeding with the final simulations, mesh convergence study was carried out to confirm stable solution is achieved and the mesh does not influence the solution. For instance, for the 8 DPH case, we built three different meshes with approximately, 90,000 cells, 140,000 cells and 300,000 cells. Mesh validation was performed by comparing peak flow speed at both inlet and outlet for each mesh cases and observed less than 1% variation between mesh 2 and 3. Then mesh with 140,000 cells was chosen for the further simulation. Irrespectively for all the DPH cases the movement of mouth cavity was simulated for 280 ms with 2,800 timesteps (each 10^−4^ s).

Flow speed at the inlet and outlet at each time step was calculated as the average of flow speed across it. Correspondingly, peak flow speed was the flow speed at the time of maximal mouth opening. Flow rates were defined as the product of flow speed and the circular area of the inlet and outlet. Peak flow rate was flow rate at the time of maximal mouth opening. Reynolds number (*Re*) was calculated as

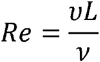

where *υ* is flow speed (m s^−1^), *ν* is the kinematic viscosity of the fluid (m^2^ s^−1^) and *L* is the characteristic length scale (m). We used the swimming speed of the larvae as the characteristic speed and the buccal length as the characteristic length.

Reynolds number was developed to characterize the flow in the case of steady flow within a long rigid tube with a fixed (time independent) radius. However, the suction flow is controlled by the rapid time-dependent motion of the cavity walls, and is characterized by strong temporal flow patterns, which needs to be considered. We therefore propose to use the Womersley number, *α*^2^ which was formulated for pulsating flows mainly associated with cardiovascular systems (Womersley, 1955), and is calculated as:

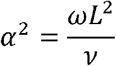

where *ω* is the characteristic angular frequency (s^−1^), *ν* is the kinematic viscosity of the fluid (m^2^ s^−1^) and L is a characteristic length scale (gape; m). The Womersley number relates the pulsation flow frequency to viscous effects. Here, the angular frequency is calculated using the time it takes the larvae to fully open its mouth (TTPG) such that *ω* = 2*π/TTPG*. Note that we used different (although correlated) characteristic lengths for the Womersley and Reynolds, referring to buccal length and gape, respectively.

## Results

To facilitate the comparison between fish species in which the growth rates can differ, we hereafter report on the scaling of suction feeding kinematics and dynamics with buccal length. As larvae mature from 8 DPH to 37 DPH, the length of the buccal cavity and the diameter of the mouth increase by about two-fold (Fig 2; Table 1). Concomitantly, the time to peak gape decreases by a factor of ^~^3.6 from an average of 57.3 ms at 8 DPH to 15.6 at 37 DPH. By and large, the kinematics observed in *S. aurata* larvae yielded unidirectional flows in our CFD models, i.e. fluid entering the mouth at the gape (inlet; Fig 5A) and exiting through the gills (outlet; Fig 5B). As previously reported (Yaniv et al., 2014), peak flow speeds at the mouth inlet (U_peak_(gape)) increased with increasing buccal length (i.e. age; *L*), following the exponential relationship U_peak_(gape) = −0.56*exp(3.39 L); (Fig 6A). Peak flow speed was 28.3 mm s^−1^ for the 8 DPH case and increased to 49.8 and 136.2 mm s^−1^ for the 23 and 37 DPH cases. Correspondingly, *Re* increases by an order of magnitude (from 23 at 8 DPH to 218 at 37 DPH).

**Fig 2.**
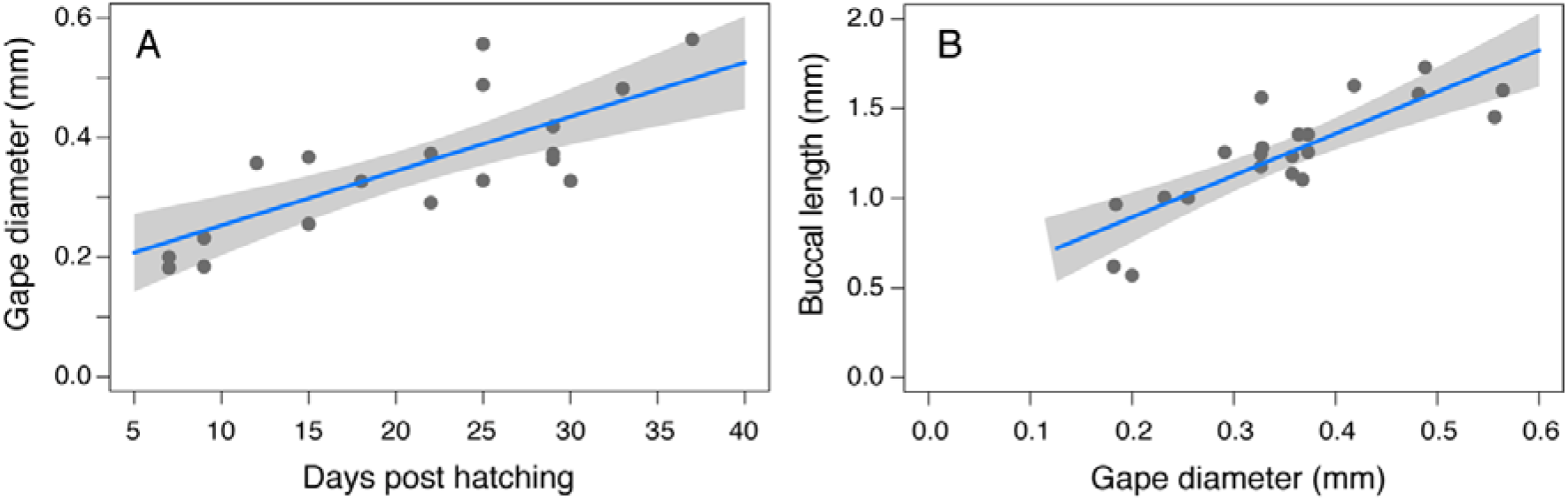
The relationships between age and gape diameter (A) and between gape diameter and buccal length (B) in *Sparus aurata* larvae ranging 8-37 DPH (n=22 individuals). Blue lines depict a linear regression between the two parameters (R^2^ = 0.55 and 0.65 for A and B respectively, P< 0.001 for both).

**Fig 3.**
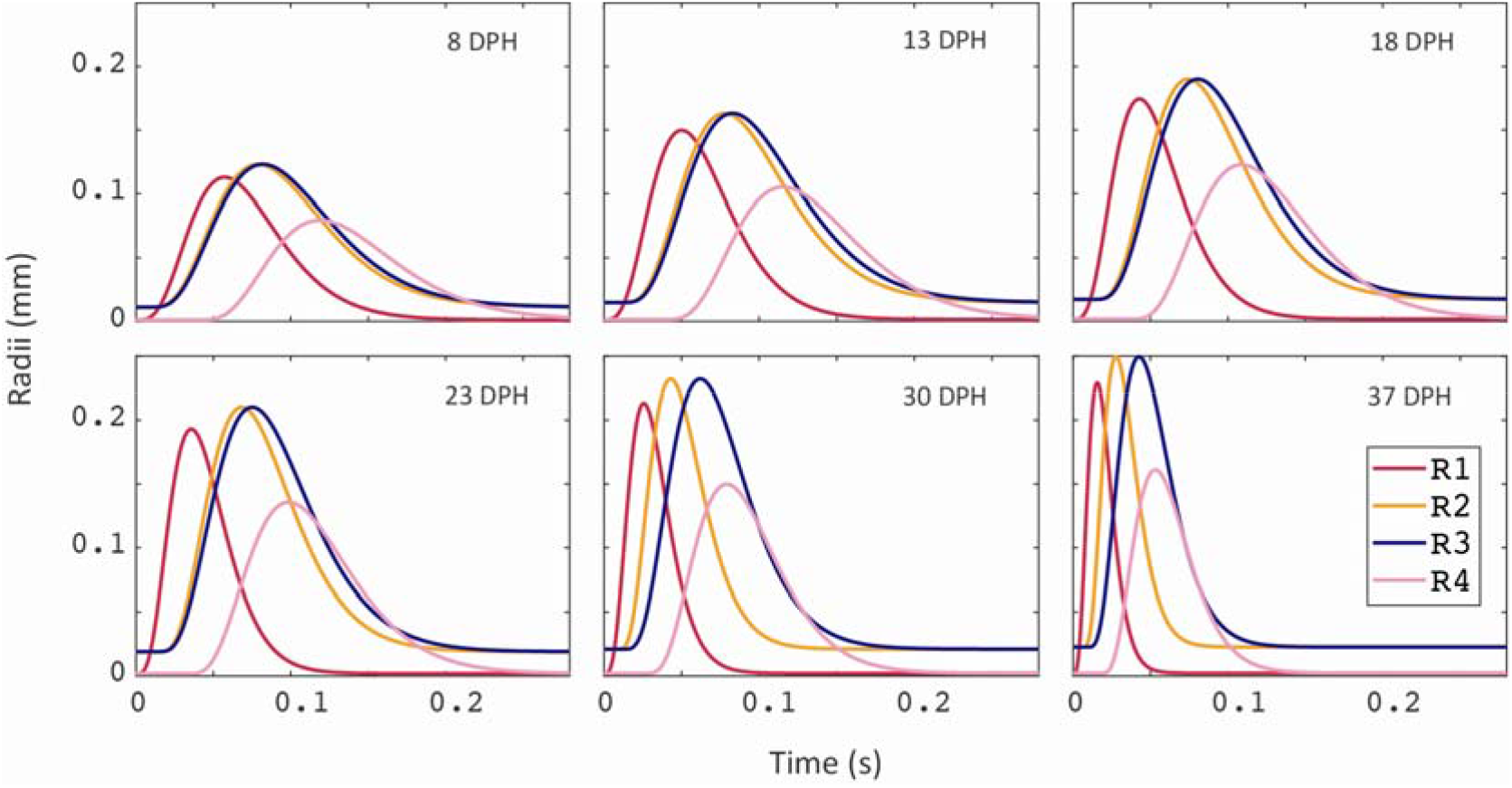
Buccal expansion kinematics across *S. aurata* ontogeny. Plots depict the radii of R_1_R_4_ as a function of time for 8, 13, 18, 23, 30 and 37 DPH larvae. Note that as larvae grow the overall time from mouth opening (R_1_) to the closing of the gills (R_4_) decreases, whereas the radii (and correspondingly buccal volume) increase. Furthermore, the timing of peak radius for each one of the mouth sections R_1_-R_4_ changes through larval growth.

**Fig 4:**
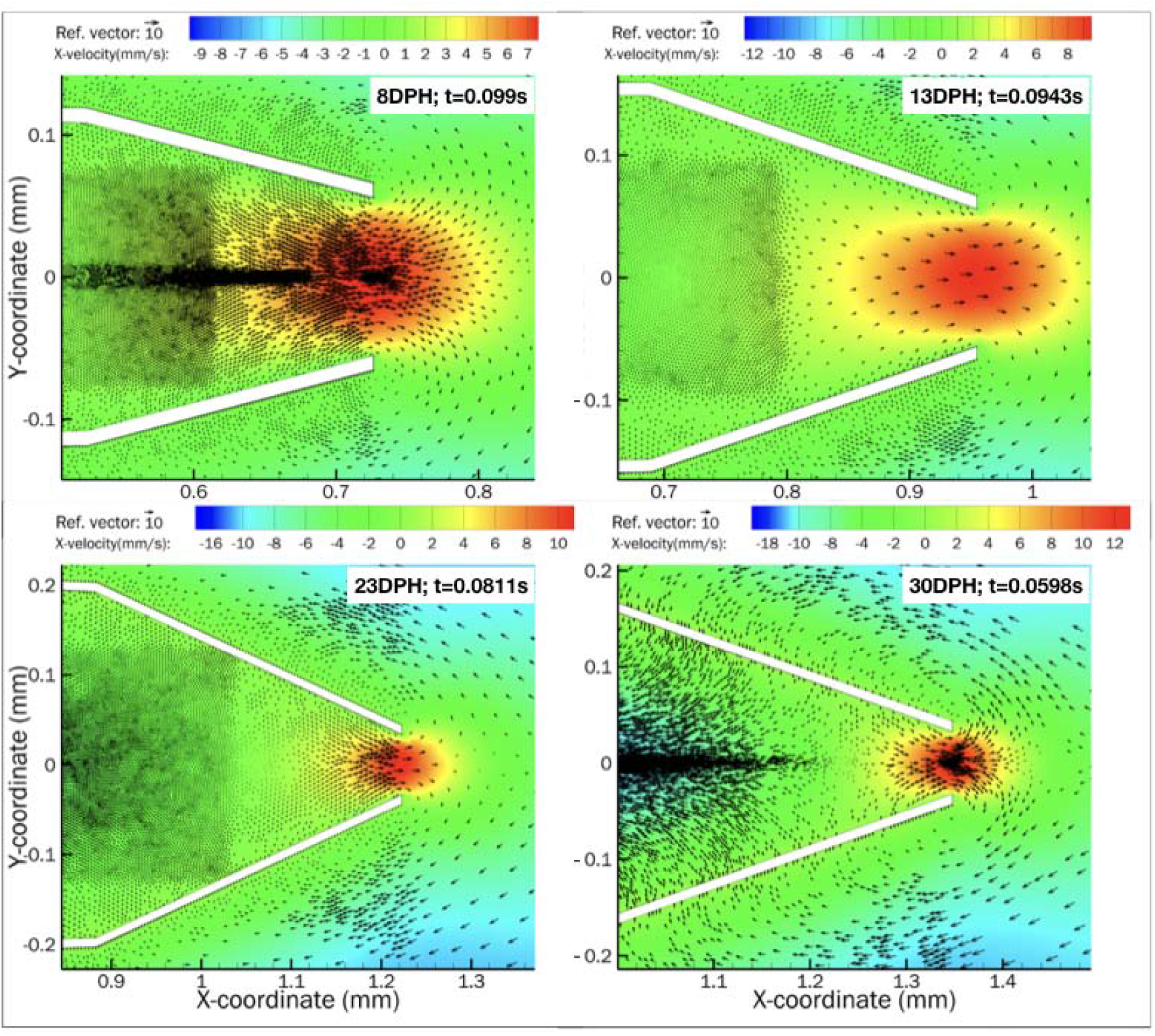
vector maps showing peak flow reversal for CFD models of 8, 13, 23, and 30 DPH larvae. Vector maps for each age were saved at the time when efflux (flux into the orifice) was maximal. Different x, y and speed scale are used in the four panels, however green color consistently represents low (and zero) flows. Also note that gape size at peak efflux decreases with increasing age.

**Fig. 5:**
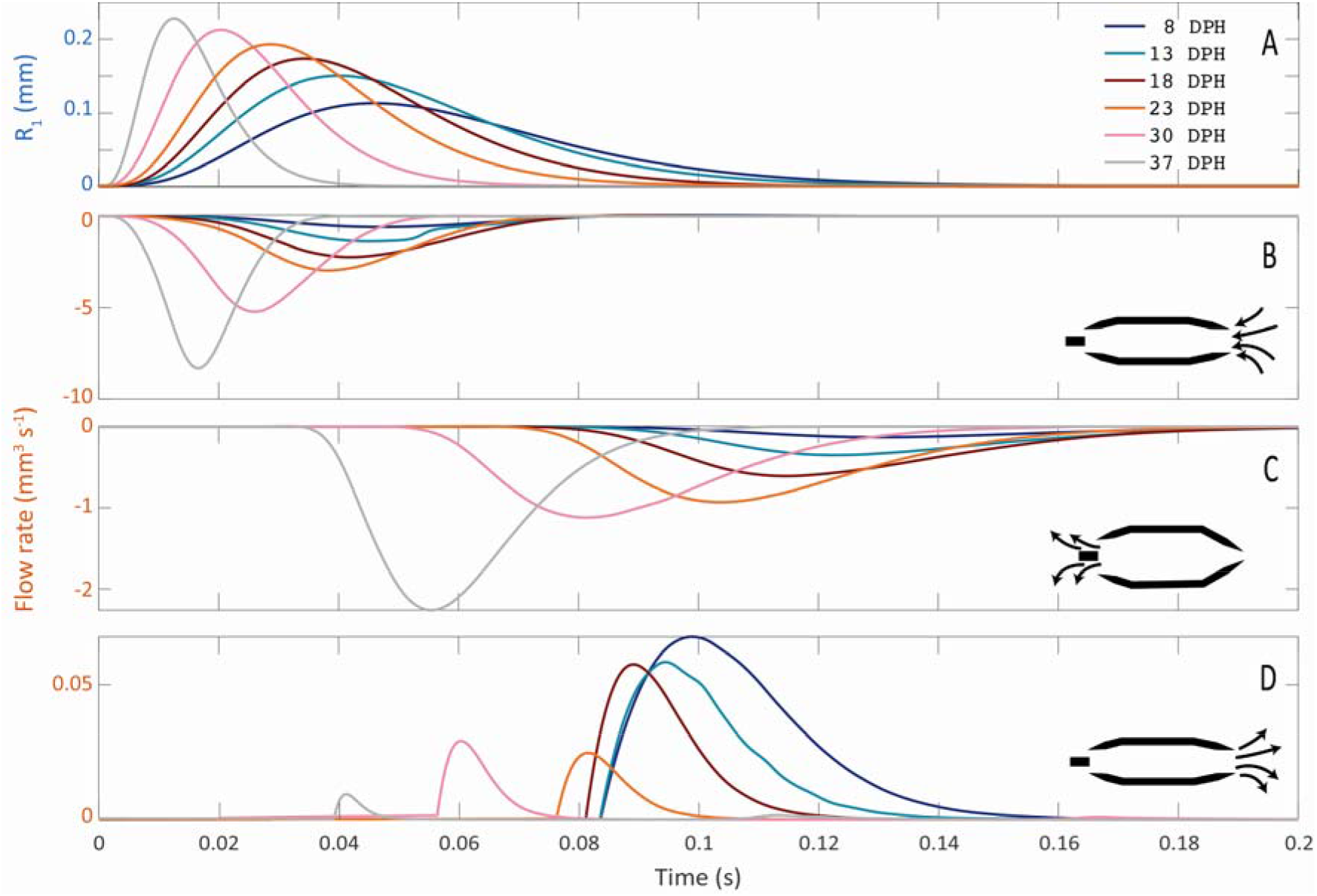
Gape size and flow rates as a function of time. Drawn are (A) the radius of R_1_ (gape size), (B) the influx into the mouth inlet (gape), (C) the efflux out of the gills and (D) the efflux out of the mouth inlet. As larvae grow, the influx at the gape and efflux at the gill increase, however the efflux at the gape (flow reversals; positive flow rate) decreases. Note the different scales and units for the Y-axes in A-D.

**Fig 6:**
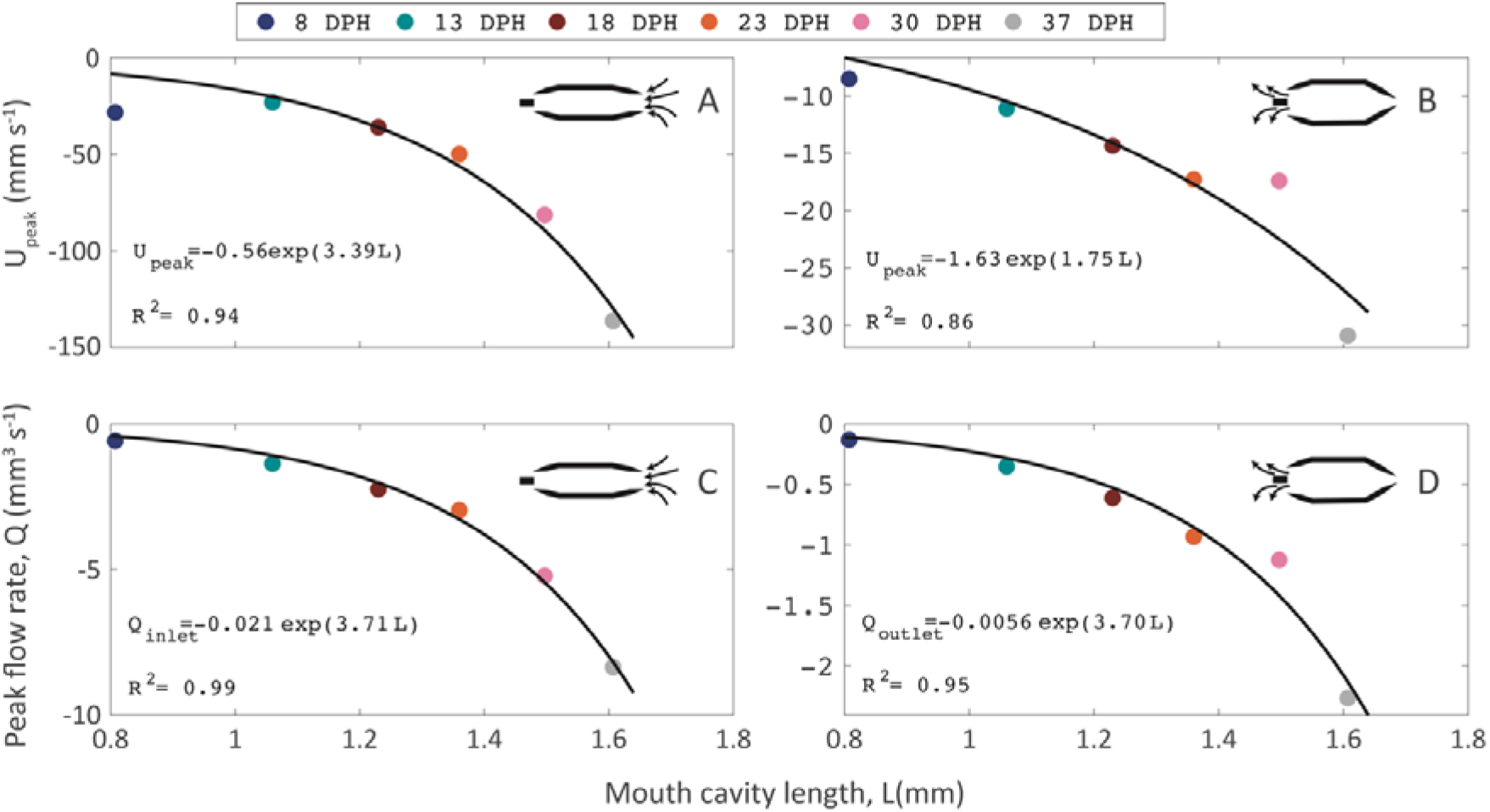
Scaling of peak flow speeds (A, B) and flow rates (C, D). Left column (A, C) depicts the inlet, right column (B, D) depicts outlet. Black lines represent exponential fits. Colors depict the different ages.

Similarly, peak flow rate at the mouth inlet (Q_peak_(gape)) increased with increasing buccal length from 0.58 mm^3^ s^−1^ for the 8 DPH case to 2.96 and 8.37 mm^3^ s^−1^ for the 23 and 37 DPH cases, following an exponential relationship Q_peak_(gape)= −0.021*exp(3.71 L) (Fig 6c). Peak flow rate at the outlet Q_peak_(gills) increased with increasing buccal length from 0.13 mm^3^ s^−1^ for the 8 DPH case to 0.93 and 2.26 mm^3^ s^−1^ for the 23 and 37 DPH cases, following an exponential relationship Q_peak_(gills)= −0.0056*exp(3.7 L).

While the observed kinematics in all cases (8-37 DPH) resulted in a net influx into the gape (inlet), we observed considerable efflux (flow reversals) around the time of mouth closure (Fig 5B). These flow reversals were most pronounced for models depicting suction feeding in young (8 and 13 DPH) ages, where efflux out of the gape was ^~^10% and ^~^3% of the influx into the cavity, respectively (Fig 6). Efflux decreased sharply for the cases of 18-37 DPH (Fig 6). Plotting the Reynolds *versus* Womersley numbers for all our cases (Fig 7) indicated that efflux at the mouth (flow reversals) was > 3% of the influx for the smaller larvae, characterized by Re < 50 and *a*^2^ < 4. Furthermore, running the model for the 23, 30 and 37 DPH cases using the observed morphology excursions but the kinematics of the 8 DPH case yielded high efflux (^~^7-8%), similar to the ones obtained for the 8 DPH case (Fig 7).

**Fig 7:**
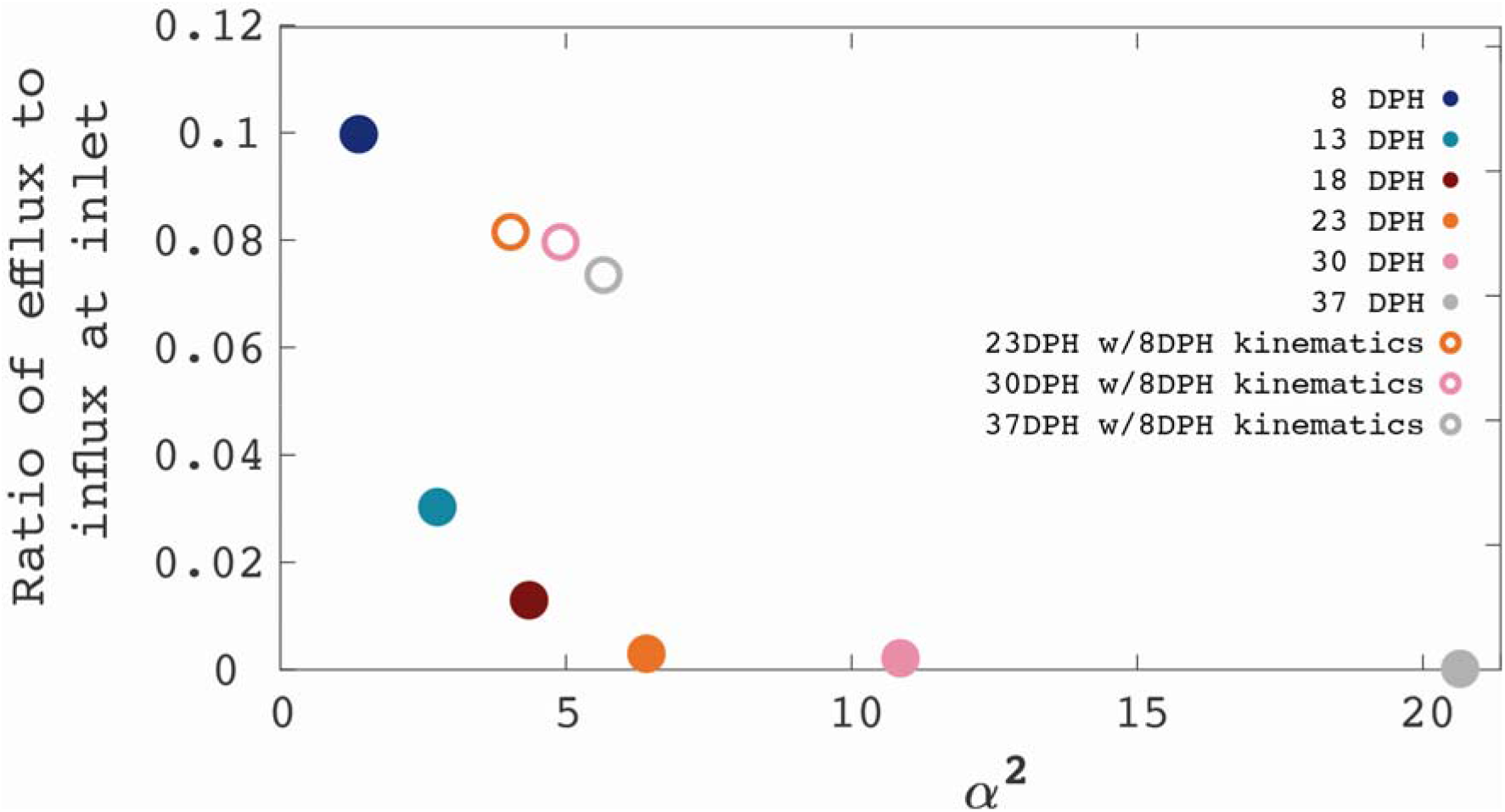
Hydrodynamic characterization of flow reversals. (A) the ratio of efflux to influx at the inlet (gape) decays as Womersley number decrease, and is most prominent at *α*^2^ < 4, indicating that flow reversals occur under conditions where viscous effects dominate over temporal (wall movements) effects. Full symbols represent the observed kinematics; open symbols represent larger models where peak excursion and time to peak excursions were similar to the 8DPH case, representing “low-effort” strikes.

## Discussion

In this study, we used computational fluid dynamics (CFD) to investigate the fluid dynamics of suction feeding larval fish. Using observed strike kinematics of *Sparus aurata* larvae ranging from first feeding to metamorphosis to parametrize the model, we quantified the flow speeds and the influx and efflux into the mouth and out of the gills for six larval ages. As larvae grow, their buccal cavity elongated and its radii increase, and it expands faster (Fig 2, Table 1). These kinematics leads to an increase in the maximal flow speed and flow rate observed at the orifice (Fig 5, 6). While most of the fluid entrained in the cavity is evacuated through the gills, we observed high efflux of water flowing outwards from the gape (Fig 5,7). These flows occurred predominantly in the models characterized by Re < 50 and *a*^2^ < 4, but also in our larger models during slow mouth opening. Overall, our results show that the inability of larval fish to capture prey may result from (at least) two hydrodynamic mechanisms: (1) their suction force do not exert sufficient force to draw the prey into the mouth (Yaniv et al., 2014), and (2) flow reversals may carry the prey outside as the mouth closes if the prey did not get deep enough into the mouth (this study; (China et al., 2017; Holzman et al., 2015)).

Previous observations of larval feeding on non-evasive prey indicate the prevalence of “in-and-out” events where prey that entered the mouth was expelled before the mouth closed (China et al., 2017; Holzman et al., 2015). The probability of these “in-and-out” events increased in suction feeding events characterized by low *Re* (<20), compared to successful events characterized by higher Re of >40 (China et al., 2017). This observation is in agreement with our results, indicating the prevalence of high efflux (flow reversals) under low Re, a condition characterizing younger larvae or older larvae that execute low-effort strikes. Furthermore, flow reversal in the models occurred later in the strike, as the mouth was closing. This timing corresponds to the observation of the “in-and-out” events, and the fact that they were initiated from a further distance compared to unsuccessful events in which the prey did not enter the mouth at all (China et al., 2017). A flow visualization study reported flow reversals in larval zebrafish, occurring in when the mouth starts closing (Pekkan et al., 2016). However, that study did not report the hydrodynamic or kinematic correlates were associated with their occurrence. For larger fish that operate at higher Re (Re >55 for 75% of >400 PIV measurements; (Jacobs and Holzman, 2018)) such flow reversals were extremely rare. In general, whether prey transport during suction feeding can hinder feeding success is rarely demonstrated.

Reynolds number is commonly used to characterize the suction flow field for adults and well as larval fishes (China and Holzman, 2014; China et al., 2017; Hernández, 2000; Holzman et al., 2015). Reynolds number provides the ratio between inertia and viscous forces; as Reynolds increase, inertia forces are considered dominated over viscous ones and vice versa. Reynolds number is frequently used to determine if the flow is laminar or turbulent (Denny and Wethey, 2001; Vogel, 1994) and for specific configurations, critical Reynolds numbers were proposed. However, given the nature of the flow within the buccal cavity, we suggest that Reynolds might not convey all the information needed to characterize the fluid phenomena. Reynolds number was developed to characterize the flow in the case of steady flow within a long rigid tube with a fixed (time independent) radius. However, the suction flow is a pressure driven flow, controlled by the rapid time-dependent motion of the cavity walls (Day et al., 2015). Hence, the boundary conditions change as the cavity opens and close over a short period of time, indicating that suction feeding is not only a pressure driven phenomena but also a transient one. Therefore, one should consider, in addition to the inertia and viscous effects, the temporal ones. Furthermore, to characterize a suction feeding event based on Re, one should choose a characteristic lengths and speed out of several possible options: for example one could justify using peak gape, or gape at the time of peak flow speed, or mean gape, and that choice would change the calculated Re. We therefore proposed to use the Womersley number, *α*^2^ which was formulated for pulsating flows mainly associated with cardiovascular systems (Womersley, 1955). While it is acknowledged that, for the case of suction feeding the repetition rate is low (i.e. unlike cardiovascular systems, the time between consecutive suction events is relatively long), the temporal parameter is dominant. Admittedly, Reynolds and Womersley numbers for different cases can be correlated because in some instances the length scales (gape and mouth length) are correlated, as well as the suction flow speed can be correlated with gape size and TTPG (Jacobs and Holzman, 2018), however we advise that future studies of suction feeding dynamics report the relevant Womersley number for their case. We re-analyzed data from (China et al., 2017) and found that failed strikes were characterized by mean *α*^2^ = 1.01 ± 0.5, “in and out” events had *α*^2^ = 1.31 ± 0.46 and successful strikes had *α*^2^ = 2.16 ± 1.33, supporting the usefulness of *α*^2^ to understand larval feeding.

We suggest that the flow reversals stem from the boundary layer that develops near the mouth’s walls, slowing the flow through the mouth. To approximately calculate the boundary layer thickness (*δ_BL_*) inside the mouth cavity and to illustrate its trend as a function of the mouth length and DPH, we utilized Blasius’s boundary layer theory for low-viscosity flow over a long plate (Falkneb and Skan, 1931). While several assumptions of the Blasius’s solution were not met in our case (e.g. steady flow over long flat plate with no pressure gradients along the flow direction), this solution should reasonably predict the trend in the thickness of the boundary layer. According to this solution, the boundary layer thickness is approximated as:

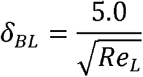

where *Re_L_*, is the Reynolds number based on the length of semi-infinite plate. In our case, we choose to estimate the boundary later thickness over the second axial length (L_2_) of the mouth cavity at an instant after the peak mouth opening where the L_2_ is almost flat and parallel to the downstream flow. We identified the time instant for the case of each DPH such as 80.4 ms, 80.4 ms, 77.5 ms, 71.7 ms, 51.5 ms, and 34.4 ms for 8, 13, 18, 23, 30, and 37 DPH, respectively. At a given time, the flow velocity of the region inside the cavity over the length, L_2_ alone was averaged and this averaged flow velocity and the length, L_2_ was used to calculate the Reynolds number (*Re_L_*). As expected, the thickness of the boundary layer decreased with increasing age. The degree of efflux exponentially increased as a function of the ratio between boundary layer thickness and gape diameter, suggesting that the development of slower flows near the cavity walls and the mouth openings could be responsible for the flow reversal.

Previous measurements (Pekkan et al., 2016), modeling (Yaniv et al., 2014) and estimations based on buccal dynamics (China et al., 2017) reported peak suction flows ranging ^~^1-40 mm s^−1^, for range of buccal length parameters (i.e. gape diameter) corresponding to the current study. While PIV measurements and CFD simulations of larval fish represent a limited sample of individuals, high-speed videos suggest that the variation in peak flow speed among individuals can be substantial (China et al., 2017). Similarly, PIV studies on adult fish indicate broad variation in peak flow speed for repeated strikes by the same individuals (Day et al., 2015; Holzman et al., 2008; Jacobs and Holzman, 2018). Such variation was not included in our study. Moreover, we base our modeling based on feeding events acquired using manually triggered high-speed cameras, and it is more likely that an observer operating it will notice and trigger an event when it is faster and greater in excursion. Thus, one should use the flow velocities estimated in our study as an example of high, rather than average, larval performance.

